# The Valuation of Social Rewards in Schizophrenia

**DOI:** 10.1101/319095

**Authors:** Lauren T. Catalano, Erin A. Heerey, James M. Gold

## Abstract

Social impairment in schizophrenia is often thought to reflect poor social cognition. Here we examine responses to social rewards, an aspect of social functioning that is not featured prominently in the literature. The goal of this experiment was to explore whether people with schizophrenia (1) undervalue social rewards, and (2) whether the undervaluation of social rewards was related to social motivation and pleasure deficits in schizophrenia and decreased social functioning. People with schizophrenia and healthy participants completed a game (Shore & Heerey, 2011) to explore preferences for different types of social (polite and genuine smiles) and nonsocial (monetary) rewards from computerized opponents. Preferences for reward types were quantified for each participant based on choice behavior during the game. Participants also completed a smile discrimination task to assess their ability to discriminate these types of smiles. Analyses revealed that people with schizophrenia (*N* = 41) treated genuine smiles as significantly less rewarding than did healthy participants (*N* = 29), despite showing a similar preference for monetary rewards. Interestingly, the undervaluation of social rewards was not related to the ability to discriminate between the smiles. The current findings provide preliminary evidence of reduced social reward valuation among individuals with schizophrenia, which may have implications for behavior in face-to-face social interactions.

**General Scientific Summary:** Social cognitive deficits are well documented in schizophrenia. However, it is less clear how people with schizophrenia respond to social rewards. Our study yields initial evidence of a deficit in social reward valuation (operationalized as a preference for genuine smiles) despite similar preferences for monetary rewards.

## The Valuation of Social Rewards in Schizophrenia

Many individuals with schizophrenia demonstrate markedly impaired social functioning. Social cognition has become a major area of interest over the past two decades because of its association with social behavior and important real-world outcomes (Green, Horan, & Lee, 2015; Couture, Penn, & Roberts, 2006; Penn, Corrigan, Bentall, Racenstein, & Newman, 1997). This literature has largely focused on the perceptual aspects of how facial expressions inform mental state inferences and impressions (For meta-analysis and review see Gur & Gur, 2015; Chan, Cheung, & Gong, 2010; Kohler, Walker, Martin, Healey, & Moberg, 2010). While this work has been informative, it remains unclear why some individuals with schizophrenia have low motivation to seek social interaction (e.g., Kalin et al., 2015; Robertson et al., 2014; Blanchard et al., 1998, 2001). Everyday functioning requires individuals to be motivated to use their social cognitive skills to navigate the social world and form relationships. There is evidence to suggest that some people with schizophrenia lack this normative social drive, which manifests as social anhedonia and/or asociality (e.g., Kalin et al., 2015; Robertson et al., 2014; Blanchard et al., 1998, 2001). Indeed, reductions in social interest and drive are evident prior to illness onset (Tarbox & Pogue-Geile, 2008; Cornblatt et al., 2012; Cannon et al., 1997) and often persist despite effective positive symptom treatment (Blanchard et al., 2001; Horan et al., 2008). A better understanding of the factors predicting motivation for social engagement is needed to elucidate the full range of social deficits associated with the illness.

Reward processing is one mechanism that drives social motivation (Depue & Morrone-Strupinsky, 2005; Ruff & Fehr, 2014). The choice to pursue social interaction is determined in part by the expectation and experience of pleasure within interaction. Social reward is often derived from social cues, such as facial displays of emotion (Blair, 2003; Bowlby, 1969). For instance, genuine smiles from a social partner elicit positive feelings that encourage future interaction, whereas a frown evokes negative feelings that motivates withdrawal to avoid danger or ill-intent. In other words, people assign value to social cues (as rewards or punishments), learn from them, and make decisions to plan for future encounters (Behrens, Hunt, & Rushworth, 2009).

It is well documented that multiple aspects of reward processing are abnormal in schizophrenia (Gold et al., 2008; Kring & Barch, 2014) and may therefore alter the expectation and/or experience of pleasure from social interaction^1^. Despite preserved hedonic capacity in nonsocial contexts (Kring & Caponigro, 2010; Kring & Elis, 2013), there may be a social-specific hedonic deficit that reduces consummatory (or “in-the-moment”) pleasure from interpersonal sources (see Cohen, Najolia, Brown, & Minor, 2010). Emerging research has yielded mixed results using different laboratory social interaction tasks (Blanchard et al., 2015; McCarthy et al., 2017); however, there is evidence to suggest that social enjoyment decreases with more severe negative symptoms (Blanchard et al., 2015; Oorschot et al., 2013; Granholm et al., 2013). Another possibility is that people with schizophrenia may not anticipate social activities to be as enjoyable (Gard et al., 2014; Engel et al., 2015), which may occur because of difficulties using social outcomes to guide behavior (Campellone et al., 2016; but also see Hanewald et al., 2017). There are few studies that directly examine reward processing of both social and nonsocial rewards (e.g., Hanewald et al., 2017).

Here we use a behavioral learning game (adapted from Shore & Heerey, 2011) to measure social and nonsocial reward valuation on the same scale. In the game, participants were asked to attempt to choose the same side of a coin as an opponent. Whenever their response matched the opponent’s response, they won, otherwise the opponent won. Our version of the task was divided into two parts. In an initial “exposure” phase, participants learned that different opponents provided different rates of monetary pay-offs (rewards on 80% of trials versus 60% of trials) and types of social feedback (genuine smiles, polite smiles, and neutral expressions). In a later “test” phase, participants chose which opponent to play from amongst pairs of opponents whose reward values they had learned. This design allowed us to quantify individuals’ preferences for reward types based on their choices during the test phase.

In this task, learning occurs gradually, as many trials are needed to develop reliable predictions of the less-than-certain outcomes. Given the evidence for relatively intact implicit reinforcement learning in schizophrenia (Heerey et al., 2008; Barch et al., 2017), we anticipated that many participants would learn the task contingencies, thereby allowing us to assess the relative preference for opponents based on the types of social and monetary feedback they provided. That is, decisions during the game should reflect reward preferences, rather than the ability to learn about the different rewards types provided. We hypothesized that because of the reduced social motivation they experience (e.g., Kalin et al., 2015; Robertson et al., 2014; Blanchard et al., 1998, 2001), people with schizophrenia would undervalue genuine smile feedback compared to healthy participants. We further hypothesized that the undervaluation of genuine smile feedback would be related to motivation and pleasure deficits and decreased social functioning in the schizophrenia group.

## Methodology

### Participants

Forty-six stabilized outpatients with schizophrenia or schizoaffective disorder (SZ) and thirty-four healthy control participants (HC) participated in the study. The SZ group was recruited from outpatient clinics at the Maryland Psychiatric Research Center (MPRC) and local community mental health clinics. Diagnosis was established using a consensus best estimate approach, combining information from past medical records, mental health clinicians, and the Structured Clinical Interview for DSM-IV Axis I Disorders (SCID-I) (First et al., 1997) based on the Diagnostic and Statistical Manual of Mental Disorders IV (DSM-IV). All SZ participants were on a stable medication regimen of constant doses and types for at least four weeks and were deemed clinically stable by their mental health clinician prior to testing. All were taking antipsychotic medications.

HC participants were recruited from the community through random digit dialing of households in nearby zip codes, word of mouth among recruited participants, and through online and newspaper advertisements. HC participants were screened with the SCID-I (First et al., 1995) and Structured Interview for DSM-III-R Personality Disorders (SIDP-R) (Pfohl et al., 1989) and were excluded if they met criteria for a psychotic disorder; bipolar disorder; and paranoid, schizotypal, or schizoid personality disorder. None of the participants met criteria for a mood episode at the time of testing. HC participants also denied family history of psychosis among their first-degree relatives and current use of psychotropic medications. All participants denied neurological injury or disease, mental retardation, as well as medical or substance use disorders within the last 6 months. Participants were between the ages of 18 and 55 years.

The Institutional Review Board at the University of Maryland School of Medicine approved the protocol (#HP-00051364). All participants provided informed consent. The SZ group completed the Evaluation to Sign Consent form (DeRenzo, Conley, & Love, 1998) to evaluate study comprehension. Compensation was $20 per hour plus a performance bonus (see below).

### Clinical Symptoms

Master’s-level raters completed the Clinical Assessment Interview for Negative Symptoms (CAINS) (Kring et al., 2013), the Brief Psychiatric Rating Scale (BPRS) (Overall & Gorman, 1962; Ventura et al., 1993), and the Calgary Depression Scale for Schizophrenia (CDSS) (Addington et al., 1992). Training consisted of an in-depth workshop on the CAINS manual and workbook by one of the developers of the instrument. A separate training covered procedures for all other measures to achieve a required reliability standard with gold-standard ratings (i.e., training videos of clinical interviews, group discussion, and instruction in interview technique). Rater agreement was not directly assessed in the current study; however, ongoing supervision was provided via regular gold-standard interview meetings to assure and maintain quality. For the CAINS, we examined the motivation and pleasure (MAP) and expression (EXP) scales (Blanchard & Cohen, 2006; Horan et al., 2011). The Specific Levels of Functioning (SLOF) (Schneider & Struening, 1983) assessed everyday and vocational functioning based on informant report. The Revised Social Anhedonia Scale (RSAS) (Eckblad et al., 1982) assessed self-reported social drive and hedonic experience.

### Neurocognition

All participants completed the Wechsler Abbreviated Scale of Intelligence—Second Edition (WASI-II; Wechsler, 1999) to assess general intelligence (4-subtest score), the Wechsler Test of Adult Reading (WTAR; Wechsler, 2001) to assess premorbid intellectual functioning, and MATRICS Consensus Cognitive Battery (Nuechterlein & Green, 2006) to assess general neuropsychological functioning. HC scored significantly higher than SZ on all measures of neurocognition (see Table 1).

**Table 1.** Participant demographic and clinical characteristics.

|  | SZ<br>(*n* = 41) | HC<br>(*n* = 29) | Statistic | *p*-value |
| --- | --- | --- | --- | --- |
| Age | 37.951 (11.872) | 39.103 (11.286) | $F(1, 68) = 0.167$ | .684 |
| Participant Education | 13.000 (2.302) | 14.621 (1.990) | $F(1, 68) = 9.398$ | .003 |
| Parental Education | 14.461 (2.540) | 13.885 (2.405) | $F(1, 62) = 0.828$ | .366 |
| Male, *n*(%) | 68.293% | 68.966% | $\chi^2 = 0.004$ | .952 |
| Race, *n*(%) | | | $\chi^2 = 0.182$ | .913 |
| African-American | 34.146% | 34.483% | -- | -- |
| Caucasian | 56.098% | 58.621% | -- | -- |
| Other | 9.756% | 6.897% | -- | -- |
| Neuropsychological Tests |  |  |  |  |
| WTAR | 100.195 (18.388) | 110.724 (9.403) | $KS\ Z = 1.525$ | .019 |
| WASI-II | 94.902 (16.550) | 108.828 (10.379) | $KS\ Z = 1.920$ | .001 |
| MATRICS | 32.220 (14.710) | 51.379 (9.049) | $KS\ Z = 2.690$ | .001 |
| Symptom Ratings |  |  |  |  |
| CAINS Total | 17.756 (8.952) | -- | -- | -- |
| MAP | 13.244 (6.426) | -- | -- | -- |
| EXP | 4.512 (3.641) | -- | -- | -- |
| CDSS Total | 1.732 (2.540) | -- | -- | -- |
| BPRS Total | 31.951 (8.282) | -- | -- | -- |
| Positive | 8.300 (4.286) | -- | -- | -- |
| Negative | 7.500 (3.289) | -- | -- | -- |
| Disorganized | 5.500 (1.895) | -- | -- | -- |
| SLOF Total | 125.610 (15.333) | -- | -- | -- |
| Interpersonal Function | 24.854 (5.655) | -- | -- | -- |
| Social Acceptability | 28.512 (1.502) | -- | -- | -- |
| Activities | 49.122 (8.340) | -- | -- | -- |
| Vocational Function | 23.122 (5.031) | -- | -- | -- |
| Chapman RSAS | 13.457 (7.139) | 8.147 (6.625) | $F(1, 68) = 12.144$ | .001 |

*Note.* WTAR = Wechsler Test of Adult Reading total score; WASI-II = 4-subtest IQ; MATRICS = composite score; CAINS = Clinical Assessment Interview for Negative Symptoms; MAP = Motivation and pleasure subscale; EXP = Expression subscale; CDSS = Calgary Depression Scale for Schizophrenia total score; BPRS = Brief Psychiatric Rating Scale; SLOF = Specific Level of Functioning Scale; Chapman RSAS - Chapman Revised Social Anhedonia Scale. Two-sample Kolmogorov-Smirnov ( $KS\ Z$ ) tests were used to compare groups on neurocognition because the homogeneity of variance assumption was violated.

### Smile Value Game

Participants played an adapted version of a “matching pennies” game (Shore & Heerey, 2011) (stimulus details are reported in the supplementary materials), designed to assess the degree to which monetary and social feedback guide learning and decision-making. In the game’s exposure phase, participants played a series of six computerized opponents, each identified by a unique face image, in a neutral/non-expressive pose. Participants’ goal in the task was to select “the same side of a coin” (heads or tails) as the opponent. “Matches” were worth 5 cents and “non-matches” 0 cents. Unbeknownst to participants, their selections did not actually matter to the reward outcomes. That is, all participants received rewards on the same number of trials, regardless of their choices. Three opponents provided match feedback on 80% of trials and the other three on 60% of trials. Two opponents (one 80%, one 60%) provided this feedback by displaying genuine smiles, two opponents (one 80%, one 60%) provided polite smile feedback^2^, and the remaining opponents retained their neutral expressions with overlaid text feedback indicating match/non-match results. Opponents who smiled on matches displayed frowns on non-match trials (neutral opponents remained neutral with a text overlay).

Participants played each opponent 25 times in random order (150 trials). Because we were unable to counterbalance reward probability, expression type, and opponent gender using only six opponents, participants only saw opponents of a single gender, counterbalanced by participant gender. Face images were counterbalanced across social/monetary contingencies.

Importantly, in this design, social and monetary feedback was fully crossed. To disentangle differences in their subjective reward values, the task includes a test phase. Test phase trials (and feedback contingencies) were identical to exposure phase trials except that participants chose which opponent to play on that trial. Participants viewed two neutrally posed opponents, presented side-by-side, and were instructed to choose the opponent they felt was “best”. Task instructions were intentionally vague so that participants were not made aware of the different win probabilities and expression types, which might have biased their choices. Once participants made their choice, the game continued as it did in the exposure phase. They saw all 15 possible opponent pairings eight times each, in random order (120 test trials). Each opponent within a given pairing appeared as often in the left position as in the right position. Opponent choices during the test phase served as the dependent variable in the task. Because participants chose between opponents in all possible pairings, we can determine the degree to which money, genuine smiles, and polite smiles contribute to choice behavior.

Finally, participants ranked the opponents from 1 (most frequently rewarded) to 6 (least frequently rewarded), without explicit instruction as to what was meant by “rewarded” (monetary or social). Participants earned five cents for every “match” trial in both the exposure and test phases and received their earnings as bonus money at the end of the study.

### Smile Discrimination Task

A smile discrimination task was conducted after the main task to assess whether SZ participants could reliably detect different smile types. In one block, participants viewed 80 static face images (40 male and 40 female), and indicated whether the expression was “neutral” or a “smile” (20 polite smiles, 20 genuine smiles, and 40 neutral expressions). In a second block, they viewed 40 images of smiling faces (20 male and 20 female), and indicated whether the face displayed a polite (20 images) or genuine smile (20 images). All tasks were presented using E-Prime (version 1.2; Psychology Software Tools). Images in both tasks were obtained from a previously validated stimulus set (Heerey, 2014). Six faces were those that had been viewed in the smile value game and the rest were novel.

### Data Analysis

#### Quantification of choice behavior

We applied the logistic response function:

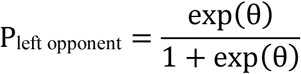

to each participant’s choice data to determine the subjective value of monetary and social feedback by examining how each feedback type contributed to a participant’s likelihood of choosing the left opponent in a given pair. In this equation, θ was modeled as:

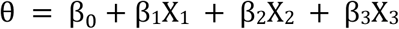

The X’s represent the differences between the left and right opponents’ monetary values (X_1_), genuine smiles (X_2_), and polite smiles (X_3_). X_1_ was coded as the difference in the opponents’ expected values (win amount multiplied by win probability). X_2_ was coded as 1 if the left opponent smiled genuinely and the right opponent was neutral, −1 if the right opponent smiled genuinely and the left opponent was neutral, and 0 if both opponents (or nether opponent) smiled genuinely. Polite smiles (X_3_) were similarly coded. The *β*s represent unstandardized regression weights for money (*β*_1_ 80% versus 60% wins), genuine smiles (relative to neutral faces; **β**_2_), polite smiles (relative to neutral faces; **β**_3_), and the intercept (**β**_0_). These values reflect the degree to which each variable contributed to choice behavior. This model allows us to quantify the effects of social rewards (genuine and polite smiles) and monetary rewards on the same scale.

The logistic regression was conducted in MATLAB (version 2015a; the Mathworks, Inc.) using an iteratively re-weighted least squares algorithm (O’Leary, 1990) to obtain the maximum likelihood estimate for each *β*. Because group-level differences in the variance of the *β*s showed violations of both the homogeneity of variance and normality assumptions necessary for the use of parametric statistical models, we examined the individually-modeled unstandardized regression weights for each model component using two-sample Kolmogorov-Smirnov tests.

#### Smile discrimination task

The smile discrimination task analysis used a signal detection theory model (Green & Swets, 1966) to code behavior. We coded a response as a “hit” if a participant correctly identified a smiling (relative to a neutral face) or a genuine smile (relative to a polite smile). “False alarms” included responses indicating the presence of a smile (or genuine smile) when the stimulus was actually neutral (or showed a polite smile). We calculated d-prime (d’) and criterion (C) for each participant (for formulas, see Macmillan & Creelman 2005). One-way ANOVAs compared average scores between groups.

## Results

### Smile Value Game

#### Responses to rewards

Data from three participants (2 HC; 1 SZ) were excluded due to a failure to follow task instructions (e.g., choosing only faces on the left in the test phase). In order to ensure that participants had learned the task contingencies (i.e., that some faces were objectively more valuable than others), we used the face-ranking data to compute an index of participants’ knowledge of reinforcement rates. On a participant-by-participant basis, we summed the rankings participants gave the 80% faces and those they gave the 60% faces and then took the difference (note that 80% faces should have received lower rankings on average than 60% faces because lower rankings are given to the most frequently rewarded opponent). There were seven participants (3 HC; 4 SZ) whose ranking difference indicated that they believed the 60% faces to be “most frequently rewarded” (difference scores of 3 or greater). This discrepancy suggested that these individuals had not understood the task. We therefore excluded these seven participants from data analysis. The final sample consisted of 70 participants (29 HC; 41 SZ; see Table 1 for sample characteristics). Excluded participants did not differ from included participants on any demographic measures or on neurocognitive or symptom measures (see Supplementary Materials). Figure 1 shows average choice behavior across different opponent pairings. As a secondary measure of learning in the task, we note that the groups did not differ in terms of their total earnings during this task (HC *M* = 9.552, *SD* = .130; SZ *M* = 9.522, *SD* = .099; F(1, 68) = 1.187*, p* =.280).

**Figure 1.**
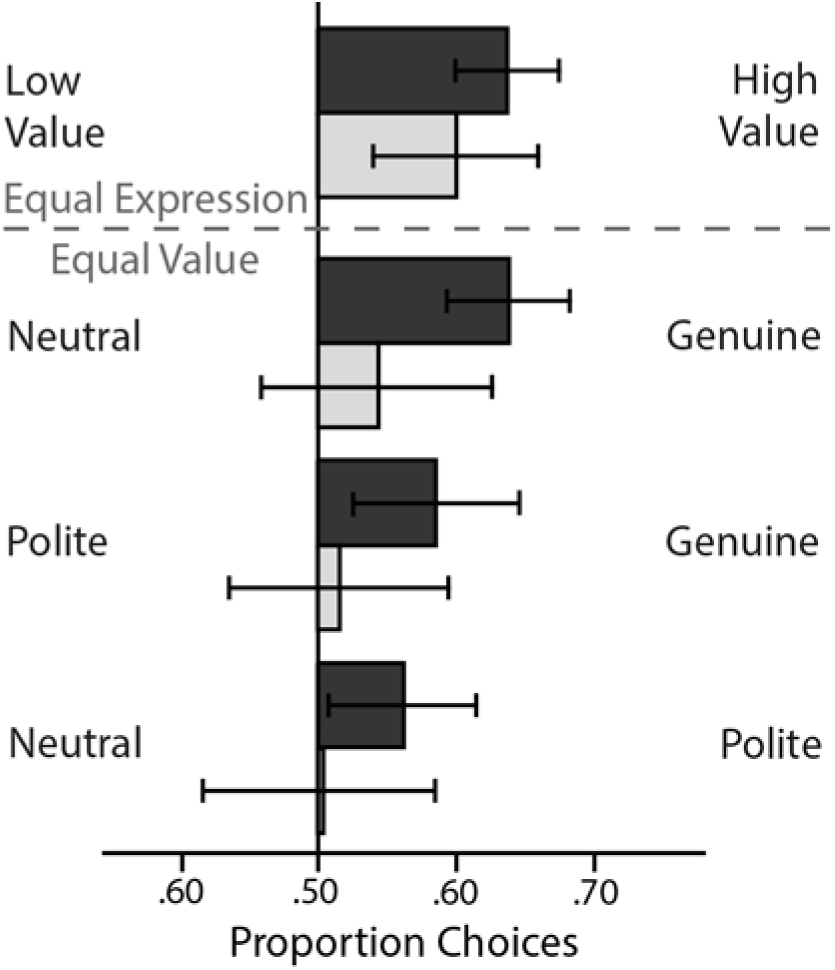
Choice behavior in the smile value game. Proportion of choices of the high-versus low-value faces when the expressions were the same (top) and proportion of choices for each expression combination when both options carried the same value (bottom). These choice data were submitted to the logistic model described above. Error bars reflect the 95% CIs.

As shown in Figure 2, HC and SZ groups did not differ in the extent to which they valued monetary rewards, two-sample Kolmogorov-Smirnov *Z* = 1.019, *p* =.250. Both groups showed clear evidence that money guided their choice behavior (i.e., median regression weights significantly greater than 0 with bootstrap-derived 95% confidence intervals that did not include 0 [see Cumming, 2014]; HC: Median: .746, 95%CI [.364, 1.128]; SZ: Median: .542, 95%CI [.293, .791]). Interestingly, HC participants valued genuine smiles (relative to neutral expressions) more than did SZ participants, two-sample Kolmogorov-Smirnov *Z* = 1.712, *p* =.006. Statistically significant two-sample Kolmogorov-Smirnov tests can be interpreted in several ways. They can mean that the two samples are drawn from non-identical distributions, that the sample medians differ, or that both of these conditions are true. Figure 2, suggests distributional differences as well as emphasizing the significant differences in the sample medians (the 95% confidence interval on the HC median does not include the SZ median, and vice versa). Thus, these data suggest reduced valuation of important social rewards in schizophrenia. Interestingly, there were no significant group differences in the degree to which polite smiles (relative to neutral expressions) guided choices, two-sample Kolmogorov-Smirnov *Z* = 1.310, *p* =.065. Genuine smiles significantly influenced HC participants’ choices, Median: 1.206, 95%CI [.417, 1.995], but did not appear to guide decisions amongst the SZ group, Median: .089, 95%CI [-.206, .384]. Polite smiles did not significantly influence choices for either group (HC: Median: .557, 95%CI [−.099, 1.213]; SZ: Median: 0, 95%CI [−.324, .324]).

**Figure 2.**
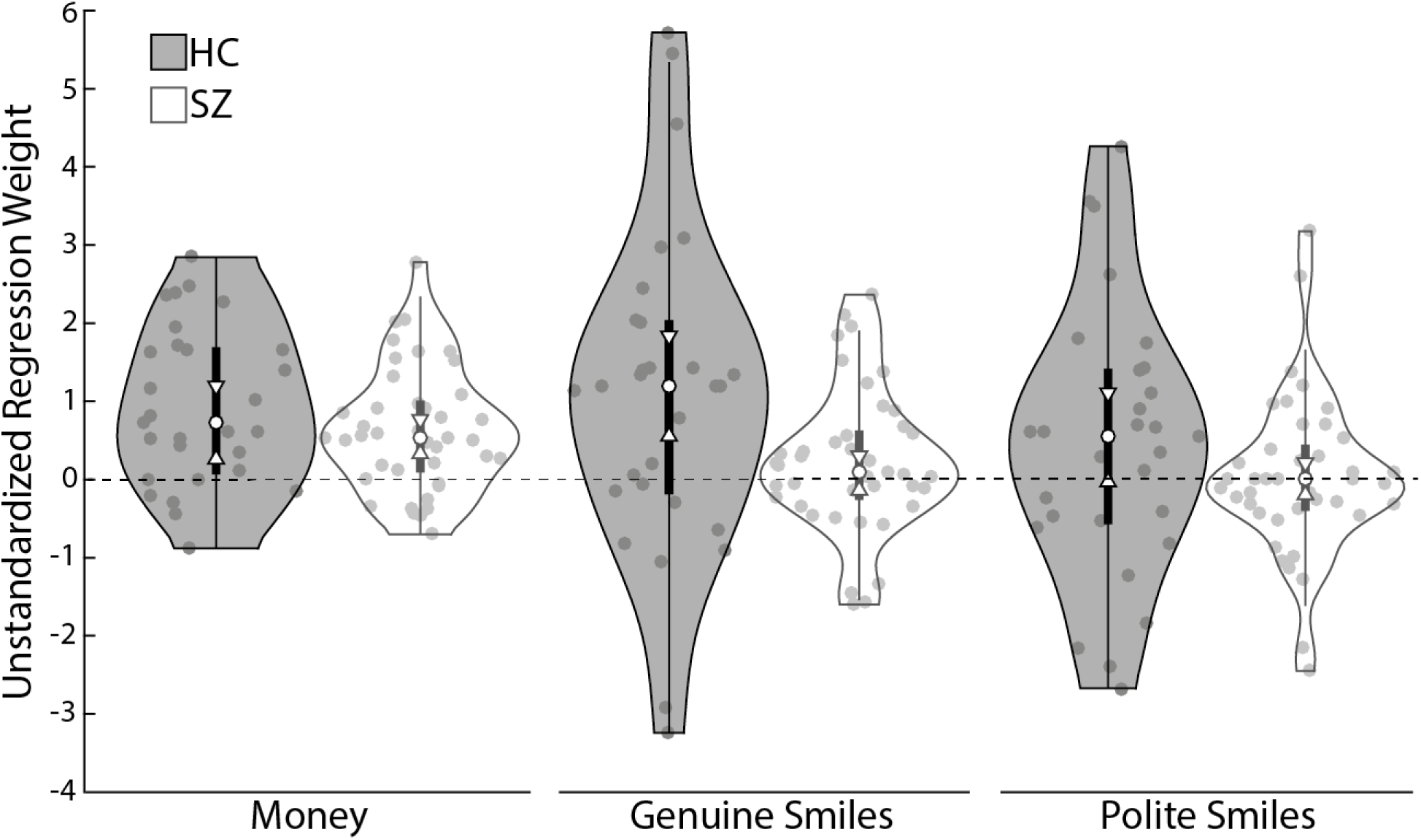
Smile value game results by group. Shaded plots show HC participants’ data and white plots show SZ participants’ data. The violin plots show the estimated probability density of the data at different values for decisions based on monetary rewards (80% opponents versus 60% opponents), genuine smiles (versus neutral faces), and polite smiles (versus neutral faces). The central shaded boxes show the inter-quartile range; the whiskers demarcate the 95^th^ percentile of the data distribution; the upper and lower boundaries on the plots show the full range of the data (including outliers); the white dots show the medians; and the white “notches” show the 95% confidence intervals on the median. Individual data points are marked with grey dots. The dashed line at zero is included as a reference point.

#### Opponent rankings

We examined participants’ rankings of the opponents descriptively, rather than statistically, because we had already used the ranking data to exclude several participants whose ranking data suggested that they had not understood task contingencies. Figure 3 shows a stacked bar plot indicating the percentage of the time that each opponent received each rank amongst participants with (hatched bars) and without schizophrenia (un-hatched bars). Qualitatively, these data suggest that more richly rewarding opponents received lower (better) rankings than less richly rewarded opponents. They also suggest that genuinely smiling opponents received lower ranks than non-genuinely smiling opponents. Finally, the data suggest that although generally similar, participants with schizophrenia may struggle to integrate social rewards into their preferences to the same degree as do those without.

**Figure 3.**
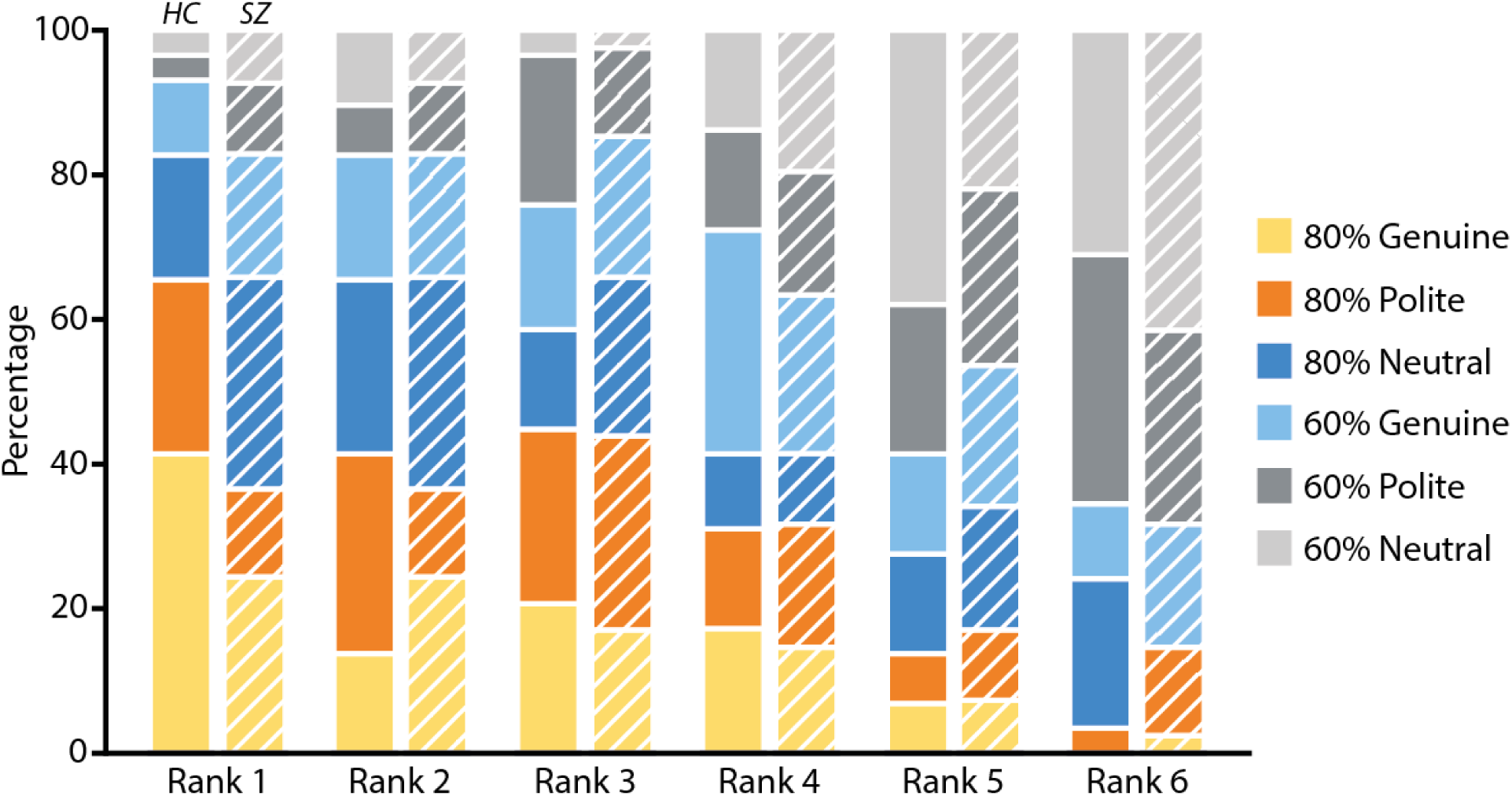
Opponent rankings. Participants explicitly ranked opponents from most frequently rewarded (Rank = 1) to least frequently rewarded (Rank = 6). Stacked bar graph shows how participants ranked each opponent as a percentage of opponents receiving each rank. Healthy control (HC) participants’ data are shown in solid bars and the data of participants with schizophrenia (SZ) appear in hatched bars. Color indicates opponent type.

#### Correlations with Clinical Symptoms, Social Functioning, and Neurocognition

We conducted Pearson’s correlations to explore how neurocognitive (HC and SZ groups) and clinical variables (SZ group) related to reward valuation (see Table 2). Note that because we are broadly interested in exploring relationships between measures, many of which we had no a priori hypotheses about, we eschew reporting traditional p-values and opt instead to show the 95% confidence intervals on the correlations. We computed the confidence intervals using a bias-corrected simple bootstrapping method (based on 10,000 samples). One can be reasonably confident that the true population statistic falls between the upper and lower bound of a 95% confidence interval. Therefore, confidence intervals that do not contain zero can be considered as reasonable evidence of an association between the two variables. Neurocognition robustly correlated with monetary preference in both groups. There were also correlations between measures of responding to genuine and polite smiles and neurocognition. Although similar, the exact patterns of smile-valuation/neurocognition correlations differed slightly across the groups. We note, however, that group comparisons of correlation strength using Fisher’s r to z transformations did not identify any significant group differences. Interestingly, within the SZ group, the valuation of genuine smiles correlated with motivation and pleasure deficits as predicted, with overall negative symptoms (CAINS-Total), and with SZ participants’ socially acceptable behavior (SLOF-Social Acceptability).

**Table 2.**
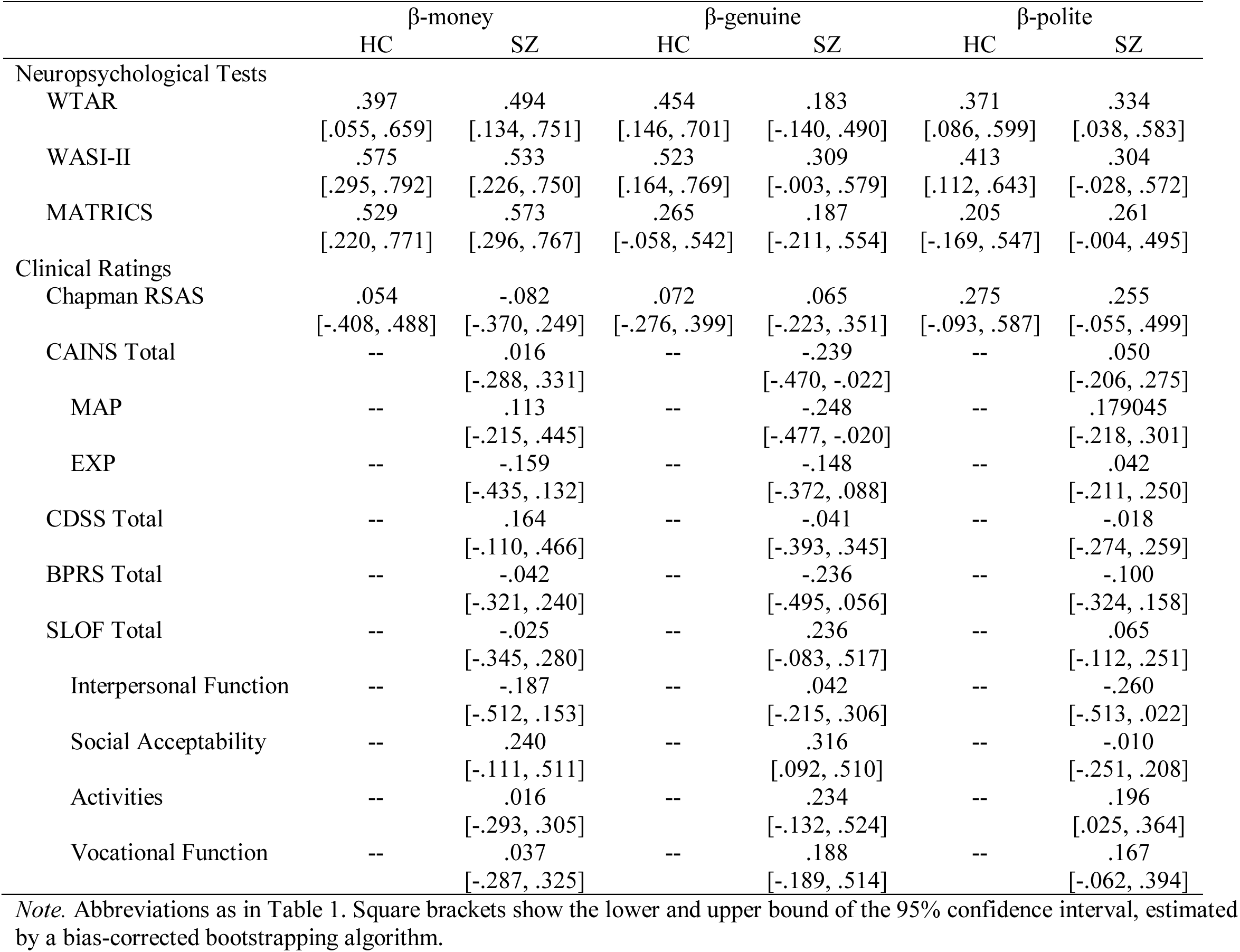
Correlations. Preferences for reward types, neurocognition, clinical symptoms, and social functioning.

| | β-money | | β-genuine | | β-polite | |
| --- | --- | --- | --- | --- | --- | --- |
|  | HC | SZ | HC | SZ | HC | SZ |
| Neuropsychological Tests |  |  |  |  |  |  |
| WTAR | .397<br>[.055, .659] | .494<br>[.134, .751] | .454<br>[.146, .701] | .183<br>[-.140, .490] | .371<br>[.086, .599] | .334<br>[.038, .583] |
| WASI-II | .575<br>[.295, .792] | .533<br>[.226, .750] | .523<br>[.164, .769] | .309<br>[-.003, .579] | .413<br>[.112, .643] | .304<br>[-.028, .572] |
| MATRICES | .529<br>[.220, .771] | .573<br>[.296, .767] | .265<br>[-.058, .542] | .187<br>[-.211, .554] | .205<br>[-.169, .547] | .261<br>[-.004, .495] |
| Clinical Ratings |  |  |  |  |  |  |
| Chapman RSAS | .054<br>[-.408, .488] | -.082<br>[-.370, .249] | .072<br>[-.276, .399] | .065<br>[-.223, .351] | .275<br>[-.093, .587] | .255<br>[-.055, .499] |
| CAINS Total | -- | .016<br>[-.288, .331] | -- | -.239<br>[-.470, -.022] | -- | .050<br>[-.206, .275] |
| MAP | -- | .113<br>[-.215, .445] | -- | -.248<br>[-.477, -.020] | -- | .179045<br>[-.218, .301] |
| EXP | -- | -.159<br>[-.435, .132] | -- | -.148<br>[-.372, .088] | -- | .042<br>[-.211, .250] |
| CDSS Total | -- | .164<br>[-.110, .466] | -- | -.041<br>[-.393, .345] | -- | -.018<br>[-.274, .259] |
| BPRS Total | -- | -.042<br>[-.321, .240] | -- | -.236<br>[-.495, .056] | -- | -.100<br>[-.324, .158] |
| SLOF Total | -- | -.025<br>[-.345, .280] | -- | .236<br>[-.083, .517] | -- | .065<br>[-.112, .251] |
| Interpersonal Function | -- | -.187<br>[-.512, .153] | -- | .042<br>[-.215, .306] | -- | -.260<br>[-.513, .022] |
| Social Acceptability | -- | .240<br>[-.111, .511] | -- | .316<br>[.092, .510] | -- | -.010<br>[-.251, .208] |
| Activities | -- | .016<br>[-.293, .305] | -- | .234<br>[-.132, .524] | -- | .196<br>[.025, .364] |
| Vocational Function | -- | .037<br>[-.287, .325] | -- | .188<br>[-.189, .514] | -- | .167<br>[-.062, .394] |

*Note.* Abbreviations as in Table 1. Square brackets show the lower and upper bound of the 95% confidence interval, estimated by a bias-corrected bootstrapping algorithm.

### Smile Discrimination Task

Groups did not differ on either smile discrimination block on measures of discrimination or response bias (*p*-values > 0.145). Furthermore, we observed no significant correlations between smile preference measures and smile discrimination performance (*p*-values > .101). This suggests that SZ participants are able to discriminate genuine from polite smiles as well as HC participants, yet do not value them to the same degree. Full descriptive data and results from the smile discrimination task can be found in the supplemental materials.

## Discussion

The present study examined social reward valuation as a mechanism that reduces social motivation in schizophrenia using a novel behavioral learning game (Shore & Heerey, 2011). Results provide two important insights. First, participants in both groups selected opponents with higher expected values to a similar degree. This finding is consistent with evidence of intact implicit reinforcement learning in schizophrenia (Heerey et al., 2008; Barch et al, 2017). Second, participants with schizophrenia failed to use genuine smiles to motivate their choices to the same extent as did healthy participants. That is, people with schizophrenia did not show the same preference to select opponents associated with genuine smile feedback. Polite smiles (a less salient social token) did not significantly shape choice behavior for either group. Although polite smiles are important social tokens, previous results suggest that they may not carry intrinsic reward value in the same way that genuine smiles do (Heerey, 2014; Heerey & Crossley, 2013). Given that the present data, as in previous research (Shore & Heerey, 2011), show that participants integrate social and monetary rewards, polite smiles may not add sufficient value to reward estimates to overcome the influence of monetary reward in these decisions. These results extend the general reward literature (Gold et al., 2008; Kring & Barch, 2014) by demonstrating that there is a unique deficit in social reward valuation (preference for genuine smiles) despite similar preferences for nonsocial rewards (money)^3^.

It would not be possible to make inferences about the relative reward value of facial expressions if participants were unable to reliably discriminate between them. Therefore, we examined whether abnormalities in social perception influenced social valuation. In the current study, the capacity to perceive smiling emotional displays was intact in schizophrenia, but these positive social cues were valued differently during the task. Importantly, smile discrimination ability did not correlate with social reward valuation. This finding is perhaps somewhat surprising in the context of the broader social cognition literature. However, our results converge with studies that show that people with schizophrenia have less difficulty identifying positive than negative emotions (e.g., Kohler et al., 2003; Mandal et al., 1998; Edwards et al., 2002; Heimberg et al., 1992)^4^. These findings provide preliminary evidence that people with schizophrenia can adequately discriminate between nuanced positive emotions (genuine and polite smiles) that are important for social affiliation, even though they may not necessarily use this information to guide their own behavior.

Why do participants with schizophrenia show specific social reward valuation deficits? One possibility is that these individuals may not learn (or might unlearn) the rewarding value of social cues because they tend to be more socially isolated and withdrawn (Oorschot et al., 2013). It is also possible that they experience less rewarding social environments because they evoke negative reactions from others due to stigma (Link, Struening, Neese-Todd, Asmussen, & Phelan, 2002) or poor social skills (Bellack, Sayers, Mueser, & Bennett, 1994). Aside from environmental factors that shape social reward valuation, a second possibility is that there are neurobiological differences that preclude individuals with schizophrenia from experiencing social feedback as rewarding (Depue & Morrone-Strupinsky, 2005; Ruff & Fehr, 2014). If this were the case, social stimuli may lose their value as the illness develops. These hypotheses are speculative and should be further explored.

Reduced valuation of genuine smiles in schizophrenia is consistent with descriptions of social anhedonia in the disorder (Blanchard et al., 1998, 2001; Horan et al., 2008). In accord, we found several associations with social-function specific clinical ratings, including motivation and pleasure. We were somewhat surprised that we did not see strong and widespread patterns emerge across more of the social, vocational and everyday functioning measures. However, it appears altogether likely that such ratings are multiply determined, reflecting both “person” variables (e.g., social interest, perception, motivation, theory of mind, etc.), as well as environment variables (e.g., ease of opportunity to engage with others, nature of living situation, availability of family members, etc.). Additionally, clinician ratings of social and vocational function may not be entirely independent of clinical symptomatology (Kalin, et al., 2015; Robertson, et al., 2014). Thus, in our view, the weak correlations with clinical rating scales, while unexpected, is not evidence of a non-relationship. Rather, the clinical endpoint may be imprecise and reflect variance from many sources. Ecological momentary assessment (EMA) might better capture real-world social behavior, providing a better endpoint to use in experimental studies. For example, EMA might provide more precise information about real-time enjoyment in social interactions, frequency of social engagement, or desire for more social contact.

Our data suggest that reduced social reward valuation, as measured by the smile value game, is a broad feature of schizophrenia that varies with the severity of negative symptoms among those with the diagnosis. That is, people with schizophrenia show reduced valuation of social rewards (i.e., genuine smiles), and this is especially true of people with schizophrenia who are more socially withdrawn and less likely to seek social interaction. A more data-driven exploratory approach in a larger sample might reveal whether social reward valuation relates to other features of the disorder (e.g., types of positive symptoms, trait negative affect, etc.).

As seen in Table 2, performance on the smile value game correlated with multiple measures of cognitive ability. This is not surprising, as one would expect measures of general intellectual ability to impact performance on a behavioral learning game. In particular, we found robust correlations between neurocognitive measures and the degree to which both participant groups valued money, in the context of similar behavioral performance on this variable. We also found similar associations with genuine smile valuation in the context of performance differences. These correlations may reflect shared variance in general underlying decision-making ability or the relationship between cognitive ability and task requirements. However, the fact that the groups showed clear differences in genuine smile valuation suggests the influence of a unique factor that may underpin in-the-moment responses to social cues. Regardless of these correlations, we suspect that undervaluation of rewarding social cues, whatever its origin, impacts social behavior and the fluidity and reciprocity present in participants’ social interactions. Future research might attempt to disentangle the impact of “general cognition” and “social cognition” by administering two separate tasks (matched for difficulty level) that differ with respect to the type of stimuli presented (social versus nonsocial). This approach might provide some ability to detect a specific deficit in social cognitive or motivational processes.

The clinical implications of these results are potentially important. Current treatments aimed at improving social functioning in schizophrenia are limited in that they narrowly focus on skill-based approaches and do not necessarily encourage individuals to seek social contact and to apply social skills outside the treatment setting (Mueser et al., 2013; also see Elis, Caponigro, & Kring, 2013). Our data suggest that social reward processing may be an important therapeutic target to foster social motivation. Future research might also examine the unique contributions of both social motivation and social cognition as they relate to social impairment. It may be the case that degraded social reward valuation early on in the course of the illness impedes social cognitive development by limiting social exposure.

Limitations of the study must be acknowledged. First, a simulated computer program served as a proxy for social interaction. Even so, social feedback altered the utility of computerized opponents, indicating that smiles are salient rewards that shape social behavior even in artificial, experimental settings. Second, social stimuli consisted solely of Caucasian, college-aged faces, which may be somewhat different to the typical interaction partner our participants, particularly those with schizophrenia, experience. Lastly, we cannot rule out the possibility that antipsychotic medications impacted performance, an important issue given the role of the dopamine system in reward processing (Salamone, et al., 2007).

## Conclusions

The idea that social cues reinforce social behavior constitutes a shift in emphasis, as most prior research in schizophrenia has focused only on social cognition. Our study yields initial evidence of reduced valuation of the social rewards that likely guide face-to-face social decision-making in schizophrenia. Additional research linking social reward valuation with the real-world use of these social cues is needed to address the degree to which reduced valuation impacts social behavior among people with schizophrenia.

## Author Note

This study was funded by the National Institute of Mental Health, grant R01 MH080066-06A1, awarded to James M. Gold. The authors have no conflicts of interest to disclose. We are grateful to the participants who helped make this study possible. We would also like to thank Sharon August, M.A. and Leeka Hubzin, M.A. who assisted with data collection.

## Footnotes

1 Basic research indicates that there is similar neural circuitry involved in nonsocial and social reward processing, with additional brain regions involved in the later (Behrens, Hunt, & Rushworth, 2009; Behrens, Hunt, Woolrich, & Rushworth, 2008; Ruff & Fehr, 2014).

2 Polite smiles differ from genuine smiles in that they can be evoked without enjoyment and do not typically activate the orbicularis oculi muscle together with the zygomaticus major muscle (Frank & Ekman, 1993; but see Krumhuber & Manstead, 2009). Evidence shows that genuine smiles carry intrinsic value over polite smiles (Shore & Heerey, 2011; Heerey & Crossley, 2013).

3 We note that these results are not related to a motivation to avoid frowns (despite the fact that some faces frowned to provide non-match feedback). Supplementary analyses of the degree to which participants avoided frowns failed to show that frowns significantly guided choice behavior in either group (see Figure S1).

4 Positive affect states may be less difficult to recognize in part because these expressions are not as complex and involve fewer facial muscles than negative emotions (Hager & Ekman, 1982), but also because they appear more frequently than negative emotions in the course of daily conversations (Fridlund, 1994).

